# Monkeypox virus M2 rewires nucleocytoplasmic transport to coordinate host translation suppression and immune evasion

**DOI:** 10.64898/2026.10.03.755596

**Authors:** Xin Huang, Yilin Yang, Wei Wang, Wuyuan Lu

**Affiliations:** Shanghai Institute of Infectious Disease and Biosecurity, Key Laboratory of Medical Molecular Virology (MOE/NHC/CAMS), School of Basic Medical Science, Fudan University, Shanghai 200032, China; State Key Laboratory of Virology, Wuhan Institute of Virology, Center for Biosafety Mega-Science, Chinese Academy of Sciences, Wuhan 430071, China; University of the Chinese Academy of Sciences, Beijing 100049, China

## Abstract

Host shutoff is a conserved viral strategy to dismantle cellular defenses, yet the molecular orchestrators in large DNA viruses remain poorly defined. Through a systematic functional screen of the monkeypox virus (MPXV) proteome, we identify M2 as a potent, multifunctional effector that executes a global blockade of host gene expression. Unlike its assembly-centric homologs across the orthopoxvirus genus, M2 localizes not only to the well-characterized endoplasmic reticulum but also to the nuclear envelope, where it may function as a molecular chokepoint, obstructing bidirectional nucleocytoplasmic transport. Concurrently, M2 expression is also associated with altered Akt/mTOR/4EBP1 signaling axis, which may contribute to reduced cap-dependent translation. Intriguingly, M2 may induce a ‘transcriptional trap’ — triggering a robust but futile ERK-dependent activation of AP-1-mediated stress responses that are decoupled from translation. Using recombinant virus engineering, we provide evidence that M2 expression augments viral replication in cell culture. These findings suggest that M2 may contribute to MPXV-mediated host shutoff, although further studies under authentic infection conditions are required to establish causality and define the underlying mechanism. Our study preliminarily reveals an paradigm for viral niche adaptation, in which a single protein coordinates spatial sequestration and biochemical suppression to enact multi-layered host suppression, providing critical candidate targets for therapeutic intervention against emerging orthopoxviruses.

## Introduction

As obligate intracellular parasites, viruses rely entirely on the host’s cellular machinery to replicate and propagate [1]. After infection, host cells initiate a series of innate immune defense mechanisms to detect viral components and trigger antiviral responses to suppress viral replication [2, 3]. In return, viruses have evolved sophisticated countermeasures to subvert these defenses, ensuring their survival and dissemination [4–6]. A central strategy employed by numerous viruses is the induction of “host shutoff”—a global suppression of host gene expression that redirects cellular resources toward viral replication [1, 7–9]. This process not only dampens antiviral signaling but also facilitates the usurpation of the translational apparatus for viral protein synthesis [10].

Host shutoff is achieved through diverse molecular mechanisms that often converge on two critical cellular processes: translational control and nucleocytoplasmic trafficking. Various factors in the mechanistic target of rapamycin (mTOR) pathway are key regulators of protein synthesis in mammalian cells, particularly the translational repressor that binds eukaryotic translation initiation factor 4E (eIF4E) or 4EBP1 [11]. Several large DNA viruses such as human cytomegalovirus (HCMV) have evolved strategies to target this translation initiation node, where mTOR complex 1 (mTORC1) is activated during early infection to maintain 4EBP1 phosphorylation; consequently, 4EBP1-sequestered eIF4E is freed up for 5’ cap-dependent translation initiation, thereby ensuring efficient viral protein synthesis [12]. Similarly, SARS-CoV-2 and picornaviruses modulate mTOR signaling to preferentially translate viral mRNAs, especially those containing internal ribosome entry sites (IRES), while repressing host translation [13, 14]. Concurrently, many viral proteins also interact with the nuclear pore complex (NPC) to disrupt its export machinery, impeding the transit of host mRNAs and proteins between the nucleus and cytoplasm [15, 16]. Notable examples include, but are not limited to, the non-structural protein 1 of SARS-CoV-2 [17, 18], the matrix protein of vesicular stomatitis virus (VSV) [19], and the infected-cell protein 27 of the herpes simplex virus type 1 (HSV-1) [20]. These coordinated attacks on translation and trafficking exemplify a convergent viral strategy to cripple host gene expression and immune effector functions.

Among the viruses capable of inducing host shutoff, poxviruses represent a particularly intriguing family due to their large DNA genomes and complex host interaction networks [21]. This is especially true for the monkeypox virus (MPXV), which caused a short-lived but dangerous global outbreak in 2022 [22]. As an enveloped double-stranded DNA virus, MPXV contains over 190 open reading frames (ORFs), and many of their encoded proteins are involved in virus–host interactions to facilitate viral replication [23]. MPXV, like other large poxviruses, stimulates immune responses more readily than smaller viruses that can rapidly replicate and disrupt host defenses [24], thus necessitating a more comprehensive strategy to survive in the host.

Vaccinia virus (VACV) is arguably the most extensively studied poxvirus, capable of hindering host protein synthesis through targeting multiple steps from pre-transcription to post-translation [25, 26]. For transcriptional control, VACV encodes two nucleases containing Nudix hydrolase motifs (D9R and D10R), which limit antiviral responses by degrading host mRNAs during poxvirus infection [27–29]. Furthermore, the viral poly(A) polymerase complex VP55/VP39 mediates aberrant hyperadenylation of host RNAs by adding a poly(A) tail at their 3’ ends, generating a group of small, nontranslated, polyadenylylated RNAs termed POLADS that sequester the poly(A)-binding protein PABP as well as impair host mRNA translation [30, 31]. At the translational level, the viral protein F17 impedes formation of the host translation initiation complex, while the viral gene VACWR169 exerts broad inhibitory effects on host protein synthesis through mechanisms yet to be elucidated [32, 33]. Additionally, VACV employs targeted protein degradation through viral factors such as C6 and C9, which mediate ubiquitin-proteasome-dependent elimination of key host antiviral proteins including HDAC4/5 and IFITs [34, 35].

Despite the strong conservation of core genes in poxviruses [36], how MPXV evades the host’s antiviral defense mechanisms remains poorly understood. Thus, we conducted a comprehensive functional screen of MPXV genes capable of globally suppressing host protein synthesis in transfected cells, leading to the identification of the M2 protein that potently inhibits host translation by disrupting the nucleocytoplasmic transport involving mRNA export and nuclear import of protein cargoes. Moreover, M2 expression correlates with suppression of cap-dependent translation and altered Akt/mTOR/4EBP1 signaling axis while concurrently activating the ERK/AP-1 transcriptional pathway, resulting in transcription but not translation of host mRNAs, a decoupling characteristic that may contribute to host shutoff. Using a recombinant VACV expressing MPXV M2, we demonstrated that M2 enhances viral virulence and promotes immune evasion. This report details how M2, transcending its ancestral role in virion morphogenesis and assembly [37, 38], contribute to host shutoff by affecting both nuclear transport and translation by strategically anchors at the nuclear envelope. However, whether these functions occur during authentic MPXV infection and whether they directly cause the observed phenotypes requires further investigation.

## Results

### Comprehensive functional screening of MPXV genome identifies multiple viral proteins as potent inhibitors of host gene expression

To systematically identify MPXV-encoded modulators of host gene expression, we constructed an expression library encompassing 164 open reading frames (ORFs) from the MPXV genome (strain Zaire-96-I-16). These ORFs, representing most of the viral proteome, were categorized into five functional groups including structural components, replication machinery, and potential virulence factors (**Fig. 1a**). Armed with this library and using the Rluc reporter gene as previously described [39], we performed a high-throughput functional screen to pinpoint viral effectors capable of suppressing host gene expression in transfected cells (**Fig. 1b)**. Briefly, HEK293T cells were co-transfected with the pRL-SV40 vector and each of the 164 plasmids encoding MPXV proteins and, after 24 h of transfection, lysed for a Renilla luciferase assay where levels of reporter gene expression were measured by fluorescence intensity or Rluc expression. As shown in **Fig. 1c**, many MPXV proteins affected reporter gene expression to varying degrees, suggesting that host shutoff may be a collaborative event coordinated by multiple viral genes of MPXV for maximum effect. Notably, a specific subset of candidates, including D10L, G7R, M2R, H5R, E4R, A10L, A12R, A18L, A19R and A35R, exhibited relatively inhibitory activity with suppression rates exceeding 85%.

**Fig. 1.**
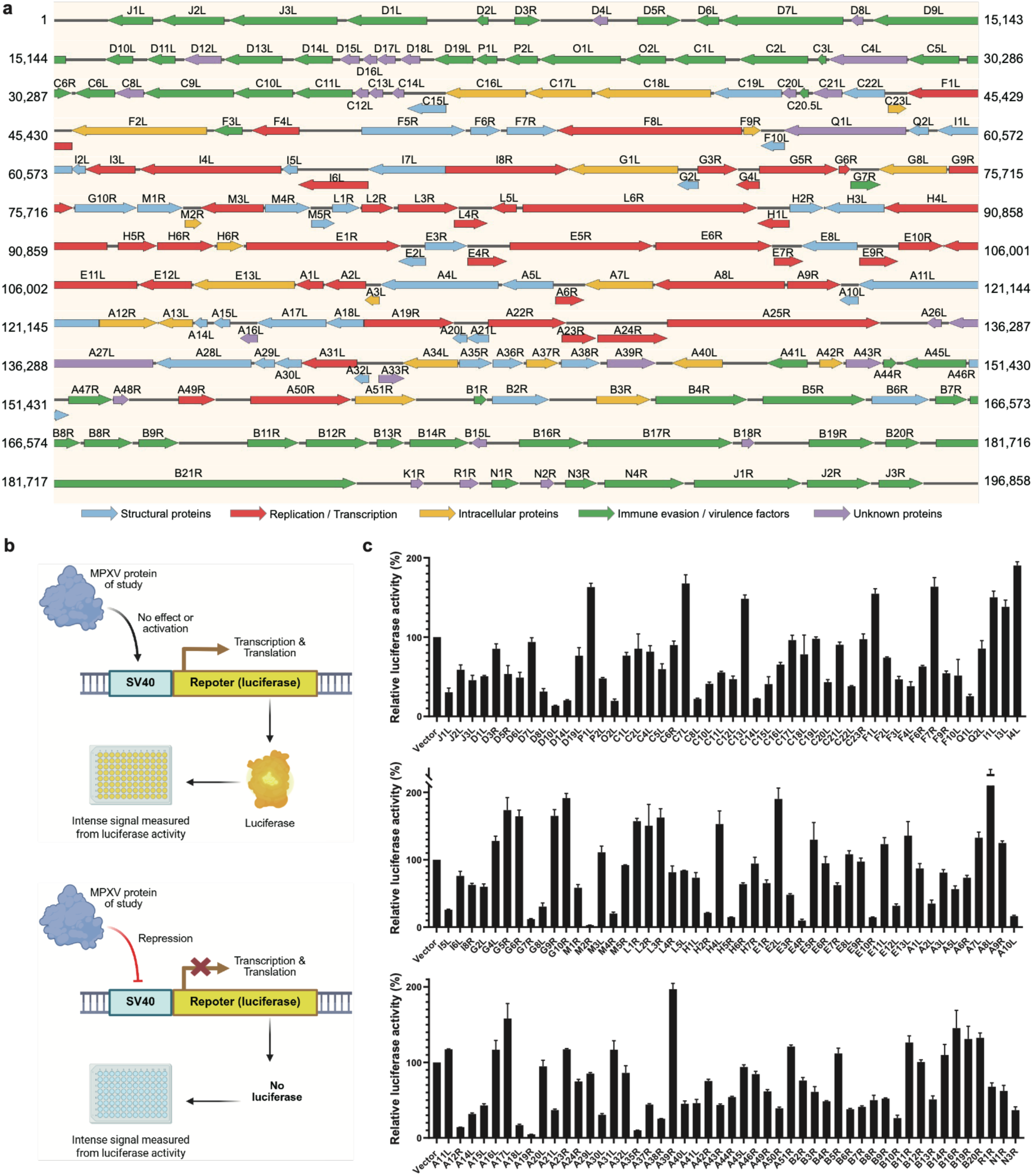
Monitor Monkeypox virus genome-induced host shutoff by preliminary screening the inhibition of Renilla luciferase (Rluc) gene expression. **a** Genome organization of the MPXV genome. The organization of ORFs on the genome of the virulent MPXV-Zaire-96-I-16 is shown. ORFs are shown as arrows to indicate their size and direction. **b** Schematic diagram of the Renilla luciferase assay screening process. c Identification of the Mpox virus genes for the ability to inhibit the Rluc gene expression. The HEK293T cells were co-transfected with pRL-SV40 and plasmids encoding MPXV viral proteins. After 24 h of transfection, the cells were lysed and detected by Renilla luciferase assay. The data are shown as mean ± SD of three biological repeats.

### The M2 protein and particularly its C-terminal transmembrane domains is the strongest driver of MPXV-mediated host shutoff among tested candidates

To validate the top candidates from our initial screen, we performed a sensitive ribopuromycylation assay, where aminoacyl-tRNA-mimicking puromycin incorporated into nascent polypeptide chains can be readily detected by Western blotting [40], to evaluate the adverse effects of the ten most potent viral genes on active translation. Among these, the nonstructural proteins M2 and A19 exhibited the most robust suppression of global protein synthesis in the assay (**Fig. 2a**), with successful expression confirmed of those ten proteins via Western blotting and immunofluorescence (**Extended Data Fig. 1a**). Notably, dose-response experiments revealed that M2 expression was associated with a more pronounced reduction in host protein abundance compared with other candidates under equivalent conditions (**Fig. 2b**). Furthermore, RT-qPCR analysis demonstrated that M2 overexpression led to a marked decrease in reporter mRNA levels (**Fig. 2c**), although we cannot exclude that this partly reflects altered RNA stability or indirect effects. Collectively, these data pinpoint M2 as a primary candidate driver of MPXV-mediated host shutoff, potentially acting at both the pre-translational and translational levels.

**Fig. 2.**
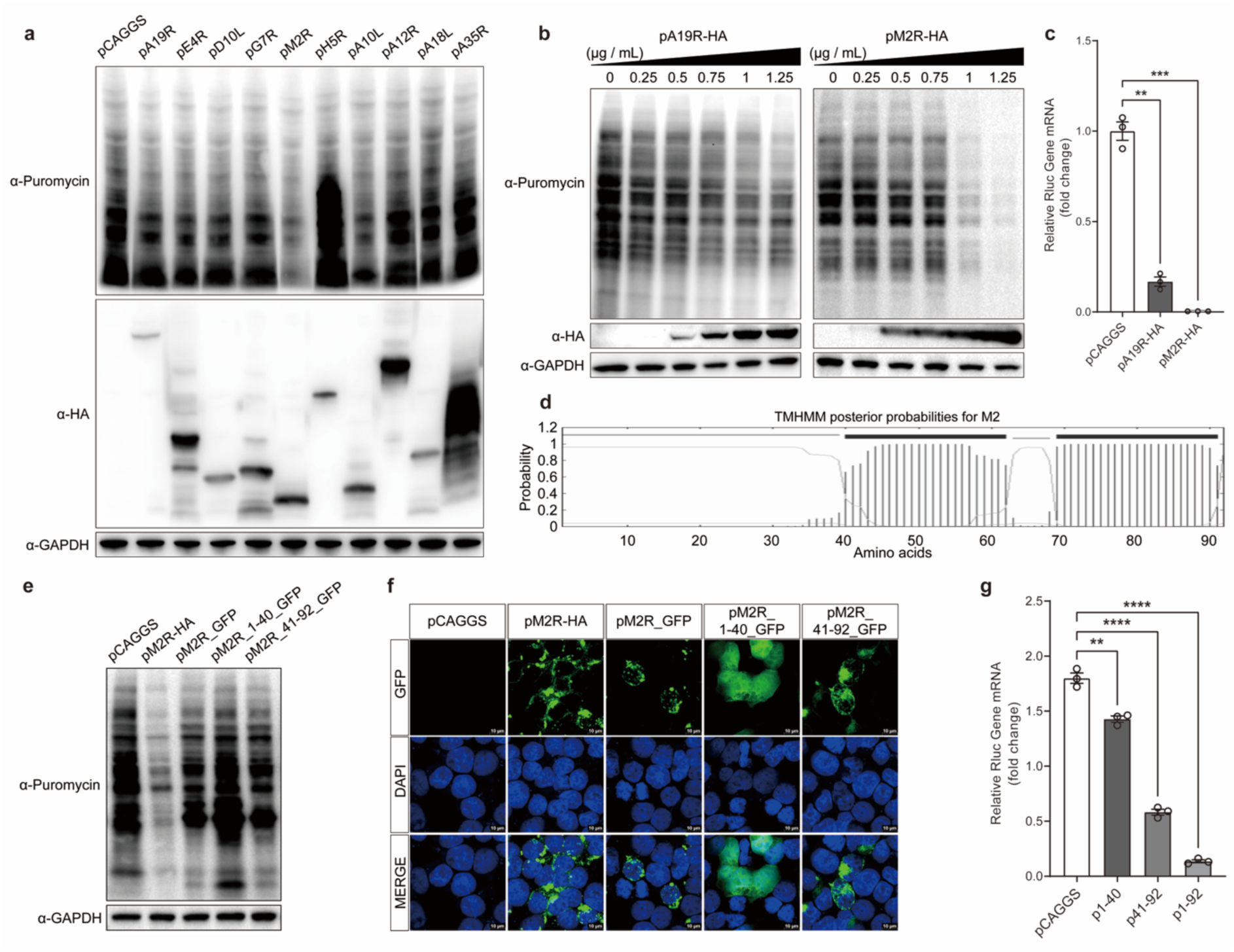
The M2 gene exhibits the strongest global translation suppression effect in host cells, with its transmembrane domain identified as the primary functional region. **a** Validation of the inhibitory effects of the top ten genes on nascent peptide chain formation at the whole-cell level. Plasmids encoding MPXV viral proteins were individually transfected into HEK293T cells. Nascent peptide chains were labeled with puromycin and detected via western blotting analysis. **b** M2 demonstrates the most significant impact on nascent peptide chain production. HEK293T cells were transfected with varying concentrations of pA19R and pM2R plasmids. Nascent peptide chains were labeled with puromycin and analyzed by western blotting. **c** HEK293T cells were co-transfected with pRL-SV40 and plasmids encoding MPXV viral proteins. After 24 hours, mRNA was extracted and subjected to RT-qPCR analysis. The data are shown as mean ± SD of three biological repeats; ***, P ≤ 0.001; **, P ≤ 0.01. **d** Prediction of transmembrane helices in M2 amino acid sequence by using TMHMM 2.0 software. The transmembrane domain is identified as the primary functional region responsible for M2-mediated global translation suppression. **e** Plasmids encoding the extracellular and transmembrane domains of M2 were constructed and individually transfected into HEK293T cells. Nascent peptide chains were labeled with puromycin and detected via western blotting analysis. **f** The expression of plasmids (pCAGGS, pM2R-HA, pM2R_GFP, pM2R_1-40_GFP and pM2R_41-92_GFP) was verified through immunofluorescence assays. **g** HEK293T cells were co-transfected with pRL-SV40 and plasmids encoding M2 fragments. After 24 hours, mRNA was extracted and subjected to RT-qPCR analysis. The data are shown as mean ± SD of three biological repeats; ****, P ≤ 0.0001; **, P ≤ 0.01.

MPXV M2 comprising 92 amino acid residues shares a high degree of sequence identity with homologs across the orthopoxvirus genus (**Extended Data Fig. 1b**). As was the case with MPXV M2 (**Fig. 2d**), they are predicated to contain an N-terminal extracellular domain (ECD) followed by two helical transmembrane domains (TMDs) at the C-terminus and implicated as a membrane-associated viral protein in virion morphogenesis and assembly [41, 42]. To delineate the main functional regions of M2, we truncated its ECD (aa 1-40) or TMDs (aa 41-92) and evaluated these constructs using ribopuromycylation and RT-qPCR assays. While both extracellular and transmembrane domains contributed to host suppression, the TMDs exhibited significantly more robust inhibitory activity than the ECD at both the protein and transcript levels (**Fig. 2e-g**). Notably, while the full-length protein remained the most potent, the TMDs appeared to be the primary functional module associated with M2-dependent host shutoff.

### MPXV M2 in place of VACV L2 augments the virulence of the vaccinia virus

To assess the physiological relevance of M2-mediated host shutoff during infection, we leveraged a VACV-based reverse genetics system to generate a recombinant virus (VM2R) wherein the L2R gene was replaced by MPXV M2 (**Fig. 3a-c**). M2 differs from L2 almost exclusively at their C-termini where the former ends with an elongated sequence, L^83^LLYTIIVLD^92^, as opposed to a shortened one ending the latter, Y^83^YFIL^87^ (**Extended Data Fig. 1b**). Remarkably, VM2R exhibited enhanced replication kinetics and significantly increased plaque formation efficiency compared to the wild-type virus (VWT) in BS-C-1 cells (**Fig. 3d, e)**. Crucially, ribopuromycylation assays during infection demonstrated that VM2R exerted a more potent suppression of *de novo* protein synthesis than VWT in a dose-dependent manner (**Fig. 3f)**, lending additional support to the importance of the C-terminal TMDs for M2-mediated host shutoff. These findings suggest that M2 could be a critical virulence factor which confers superior host-manipulation capabilities to the virus within an authentic infection landscape.

**Fig. 3.**
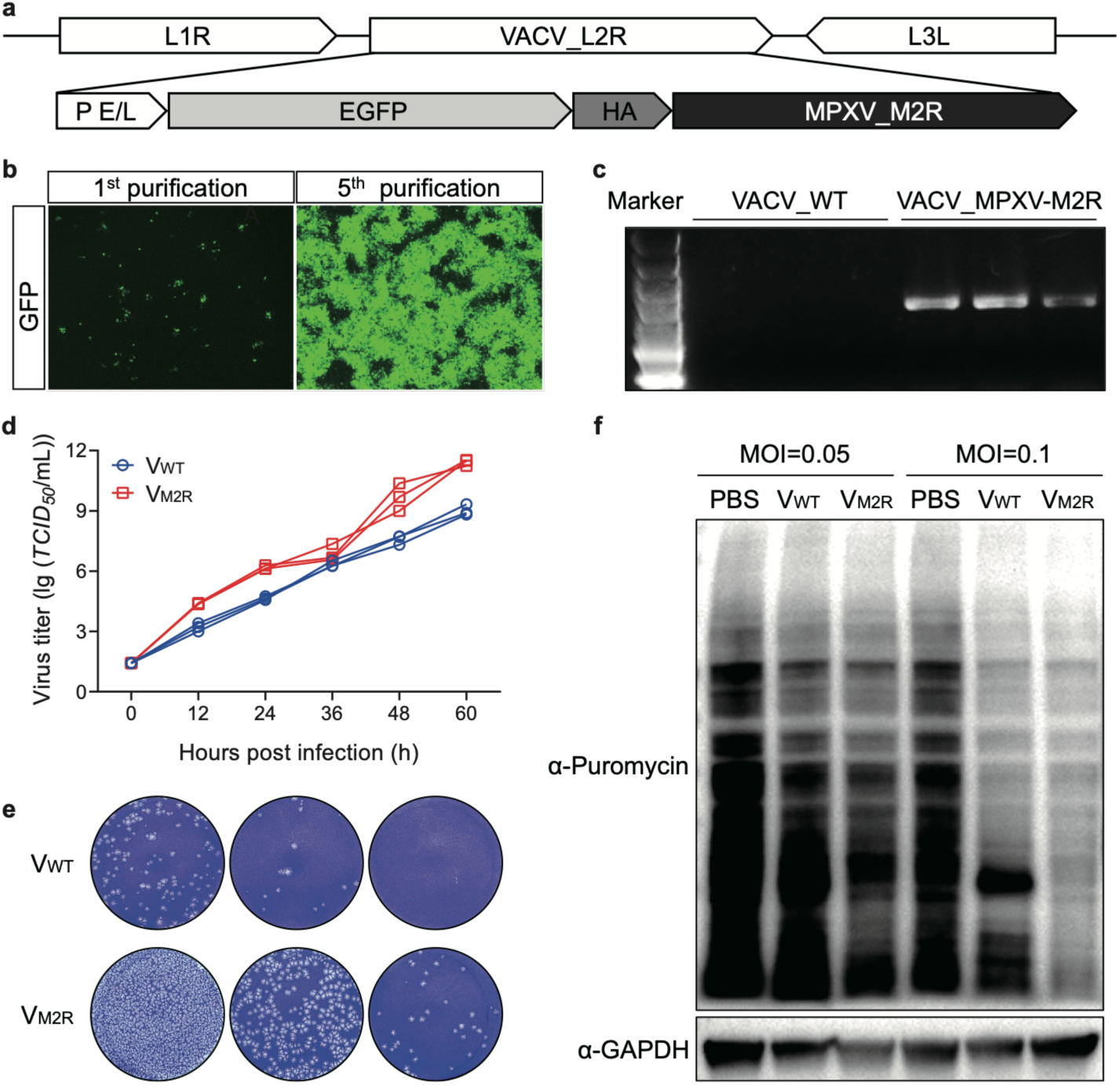
Construction and characterization of a recombinant vaccinia virus (VACV) expressing HA-epitope-tagged MPXV M2. **a** Schematic diagram of the VM2R genomic architecture. P E/L, a VACV late promoter; EGFP, enhanced green fluorescent protein open reading frame; HA, influenza virus hemagglutinin epitope tag. Verification of M2R gene expression in recombinant virus. **b** The expression of fluorescence in the recombinant virus was observed after the first and fifth rounds of purification. **c** PCR experiments were conducted to detect the expression of the inserted target gene in both the wild-type and recombinant viruses. **d** Viral growth kinetics. BS-C-1 cells were infected with VWT or VM2R at a multiplicity of infection (MOI) of 0.01. At 0-, 12-, 24-, 36-, 48-, and 60-hour post-infection, cells were harvested, and viral titers were determined by TCID50 assay. **e** Viral virulence assay. BS-C-1 cells were infected with serial dilutions of the recombinant virus VM2R or wild-type VACV WR (VWT). After 48 hours, cells were fixed and stained with crystal violet. **f** Effect of recombinant viruses on global cellular translation. BS-C-1 cells were infected with two different dilutions of VM2R or VWT. After 24 hours, nascent peptide chains were labeled with puromycin and detected by western blot analysis.

### M2 partially localizes to the nuclear envelope to exert host-shutoff function

Given that M2’s VACV homolog, L2, is an ER-resident protein [38], we examined M2’s subcellular localization using immunofluorescence microscopy. Beyond its expected ER presence, M2 also partitioned to the nuclear envelope (**Fig. 4a**). Although M2 overexpression induced ER swelling as shown by transmission electron microscopy (**Extended Data Fig. 2a**), it failed to trigger a canonical ER stress response characterized by elevated levels of stress-related factors such as PERK, BIP, and ATF4 (**Extended Data Fig. 2b, c**). While the expression of eIF2α at the transcription level was slightly increased by M2, thare was no significant change in the protein synthesis level. These data suggest that M2-mediated shutoff might be independent of ER stress-induced translational arrest. Importantly, time-course analysis revealed that the onset of global translation inhibition started at 18 h post-transfection, largely coinciding with M2’s partially translocation to the nuclear fraction (**Fig. 4b, c**). At 12 h, M2 was predominantly restricted to the cytoplasm with no detectable host suppression; however, by 24 h, its pronounced nuclear envelope accumulation mirrored the dramatic reduction in host protein synthesis. These findings establish a strong spatiotemporal correlation between M2’s nuclear localization and its potent host-shutoff activity.

**Fig. 4.**
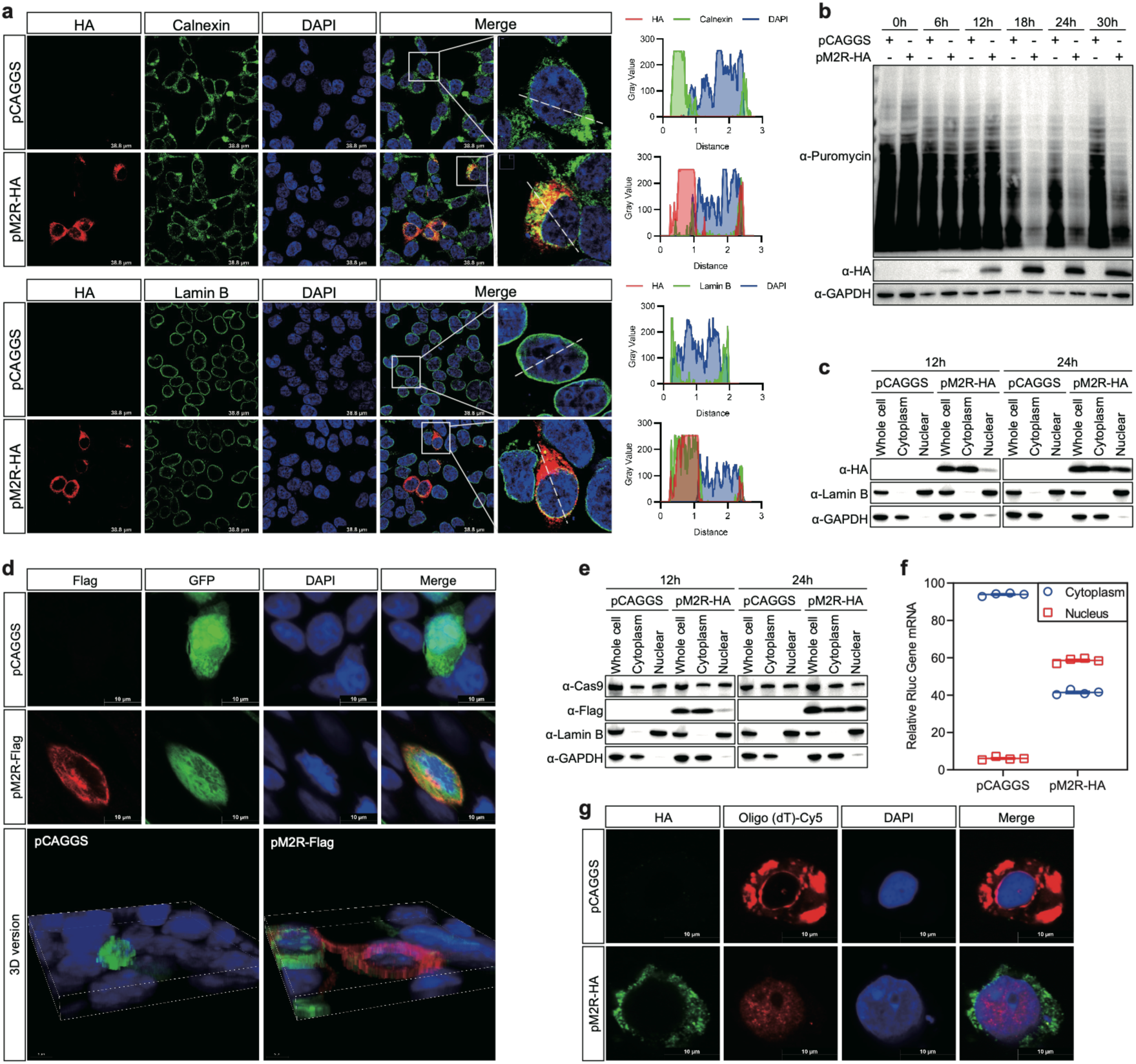
The global translation shutdown induced by the M2 protein is positively correlated with its localization at the nuclear envelope and M2 impairs nucleocytoplasmic transport. **a** Localization analysis of M2 by immunofluorescence. HEK293T cells transfected with pCAGGS and pM2R-HA were fixed and permeabilized at 24 hours post-transfection. Cells were co-stained with rabbit anti-calnexin polyclonal antibody (ER marker) or rabbit anti-lamin B polyclonal antibody (nuclear envelope marker), along with mouse anti-HA monoclonal antibody to detect M2. Secondary antibodies used were goat anti-rabbit IgG conjugated to Alexa Fluor 488 and goat anti-mouse IgG conjugated to Alexa Fluor 594, respectively. Nuclei were counterstained with DAPI. **b** Time-dependent effect of M2 on global translation shutoff. HEK293T cells were transfected with pCAGGS and pM2R-HA. Nascent peptide chains were labeled with puromycin at 0, 6, 12-, 18-, 24-, and 30-hour post-transfection. Cell lysates were analyzed by western blotting using anti-puromycin and anti-HA antibodies, with anti-glyceraldehyde phosphate dehydrogenase (GAPDH) antibody serving as the loading control. **c** Subcellular localization of M2 assessed by nuclear-cytoplasmic fractionation. HEK293T cells transfected with pCAGGS and pM2R-HA were harvested at 12- and 24-hours post-transfection. Nuclear and cytoplasmic fractions were separated and analyzed by western blot using antibodies against the HA tag, the endoplasmic reticulum marker BIP, the nuclear envelope marker lamin B, and GAPDH as a loading control. M2 inhibits nuclear protein import. HEK293T cells were co-transfected with pCAGGS or pM2R-Flag together with pMJ920 (expressing a nuclear-localized protein). **d** After 24 hours, cells were immunostained with an anti-Flag primary antibody followed by a DyLight 594-conjugated secondary antibody and visualized by fluorescence microscopy. **e** Alternatively, transfected cells were subjected to nuclear-cytoplasmic fractionation, and the distribution of the target protein in each fraction was analyzed by western blotting. (f and g) M2 inhibits nuclear mRNA export. **f** HEK293T cells were co-transfected with pCAGGS or pM2R-HA along with pRL-SV40. After 24 hours, nuclear and cytoplasmic fractions were isolated, and the relative distribution of Renilla luciferase mRNA in each compartment was quantified by RT-qPCR. **g** HEK293T cells were co-transfected with pCAGGS or pM2R-HA. At 24 hours post-transfection, cellular mRNA was labeled using a Cy5-conjugated oligo(dT) probe via fluorescence in situ hybridization (FISH), while M2-HA was detected using an anti-HA primary antibody and a DyLight 488-conjugated secondary antibody.

### M2 disrupts bidirectional nucleocytoplasmic transport

Given M2’s partial nuclear envelope residency, we investigated its impact on nucleocytoplasmic trafficking. Using NLS-tagged Cas9-EGFP that efficiently transports to the nucleus as a reporter [43], we observed that M2 expression caused significant cytoplasmic retention of the cargo, as confirmed by both immunofluorescence and subcellular fractionation (**Fig. 4d, e**). Conversely, FISH analysis using oligo(dT) probes revealed a dramatic nuclear sequestration of poly(A)+ mRNA in M2-expressing cells, and RT-qPCR of nuclear and cytoplasmic fractions showed a significantly increased nuclear proportion of Rluc mRNA transcribed from pRL-SV40, potentially indicating a concurrent block of mRNA export (**Fig. 4f, g**). To further dissect the stage of inhibition, we employed 5-ethynyluridine (EU) labeling and mRNA transfection assays, which ruled out global transcriptional suppression and direct mRNA degradation (**Extended Data Fig. 3, 4**). Polysome profiling further corroborated a shift from active polysomes to 80S monosomes, consistent with a depleted cytoplasmic mRNA pool (**Extended Data Fig. 5**). Collectively, these data suggest that M2 might be a potent disruptor of bidirectional nucleocytoplasmic transport, potentially decoupling host transcription from translation.

### M2 modulates host translation via the mTOR/4EBP1 Pathway

To explore the biochemical basis of M2-mediated host shutoff, we performed quantitative proteomics, which highlighted changes in multiple pathways with lipid metabolism being among the most prominent, and pathway analysis also suggested a significant enrichment of the mTOR signaling axis (**Fig. 5a-f**). Immunoblotting showed that M2 expression was associated with dose-dependent dephosphorylation of Akt and its downstream effector 4EBP1, while other components like eIF4G and P70S6K remained relatively stable (**Fig. 5g, h**). Notably, the marked reduction in p-4EBP1—the active repressor of cap-dependent initiation— coincided with a total loss of Renilla luciferase reporter expression (**Fig. 5i**). These results are consistent with the model in which M2 orchestrates host shutoff is associated with systematically altered the Akt/mTOR/4EBP1 signaling axis to dismantle the cap-dependent translational machinery.

**Fig. 5.**
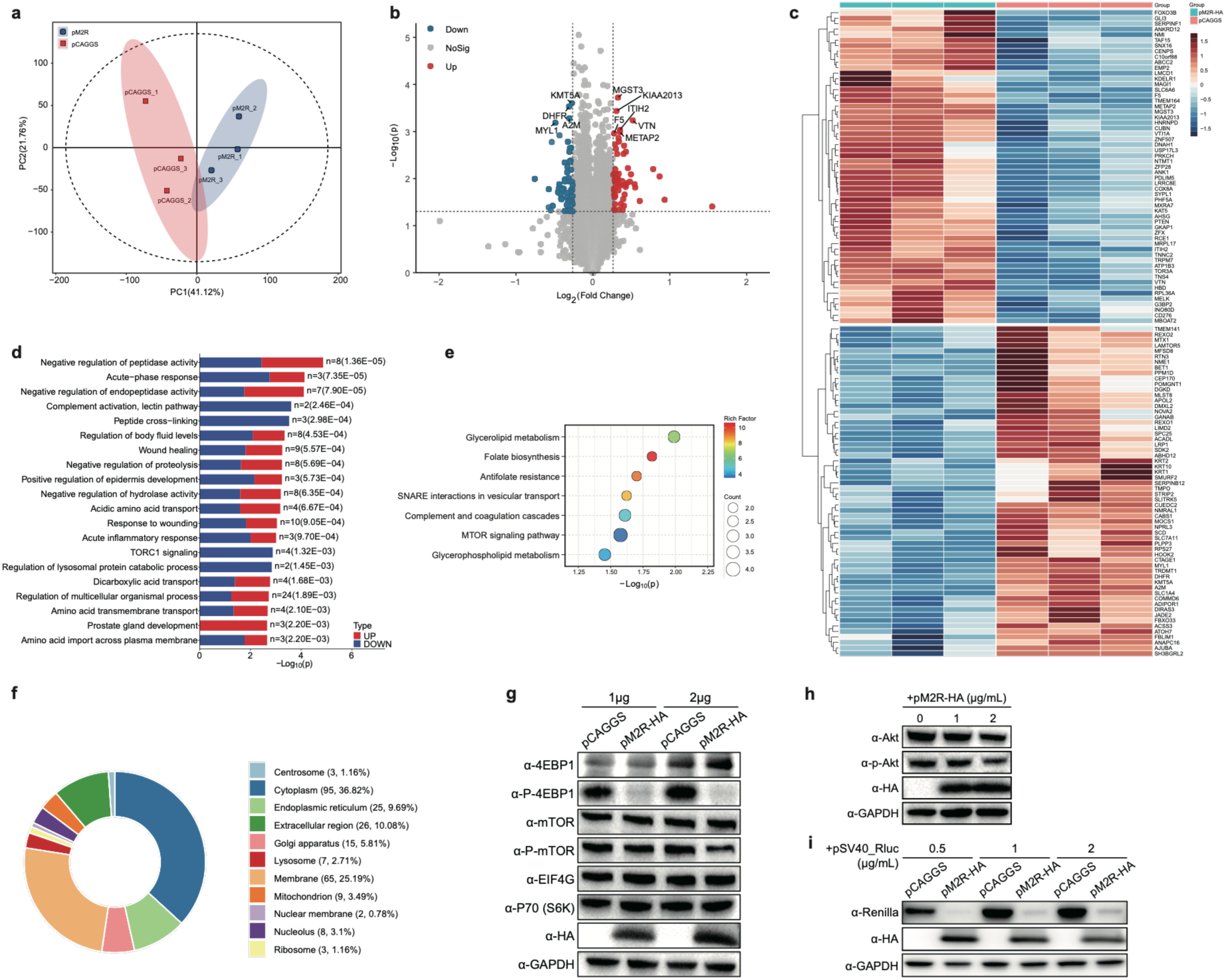
M2 modulates cap-dependent translation through the mTOR/4EBP1 pathway. **a** Principal Component Analysis (PCA) of proteomic data showing clear separation between M2-expressing (red) and control (blue) samples, indicating distinct protein expression profiles. **b** Volcano plot represents the differential expression of proteins between M2 and control groups. Red and green points indicate significantly upregulated and downregulated proteins, respectively, with a p-value threshold of 0.05. **c** Heatmap showing the hierarchical clustering of the top differentially expressed proteins, highlighting distinct expression patterns between M2-expressing and control cells. **d** Bar chart of enriched biological processes and cellular pathways, highlighting the most significantly altered functions in M2-expressing cells. Key enriched terms include negative regulation of peptidase activity, wound healing, and TORC1 signaling. **e** Bubble plot displaying enriched Kyoto Encyclopedia of Genes and Genomes (KEGG) pathways, with the mTOR signaling pathway identified as a prominent pathway significantly altered by M2 expression. **f** Subcellular localization distribution of differentially expressed proteins, showing a majority in the cytoplasm and endoplasmic reticulum, with a small fraction localizing to mitochondria and lysosomes. **g** Analysis of the effect of M2 on the mTOR/4EBP1 signaling pathway. HEK293T cells were transfected with increasing concentrations of pM2R-HA or empty pCAGGS vector control for 24 hours. Proteins from whole-cell extracts were analyzed by western blotting using antibodies against the indicated signaling molecules. **h** Examination of M2 effects on Akt activation. HEK293T cells were transfected with graded concentrations of pM2R-HA. At 24 hours post-transfection, cell lysates were subjected to western blot analysis. **i** Validation of M2-mediated suppression of cap-dependent translation. HEK293T cells were co-transfected with varying concentrations of pRL-SV40 and pM2R-HA. After 24 hours, cells were harvested and lysed, followed by western blot analysis.

### M2 reprograms host cell transcription through ERK-dependent activation of AP-1

Finally, we integrated transcriptomic (RNA-seq) and proteomic (IP-MS) analyses to explore the strategic logic of M2-mediated host shutoff. While M2 induces a robust transcriptional ‘alarm’—evidenced by ERK-dependent activation of the AP-1 family (FOS, JUN) (**Fig. 6a-f**)— this response is rendered immunologically silent. IP-MS and structural modeling identified the RNA helicase DDX51 as a candidate interactor of M2 (**Fig. 6g-i**). Given DDX51’s reported essential role in RNA splicing and nucleocytoplasmic transport, its potential sequestration by M2 likely acts as the molecular ‘chokepoint’. This strategic blockade might create a profound decoupling of host cell signaling: the cell would vigorously transcribes antiviral and stress-related messages that are subsequently trapped within the nucleus or excluded from the translational machinery (**Fig. 4f-g, Extended Data Fig. 5b**). By potentially affecting both nuclear pore function and mTOR-dependent translation initiation, M2 may ensure that despite a robust transcriptional response, the host fails to mount an effective protein-level defense, thereby facilitating unabated viral dominance. Taken together, our study identifies M2 could be a master regulator that orchestrates a dual-layered host shutoff, decoupling transcription from translation to ensure viral immune escape (**Fig. 7**).

**Fig. 6.**
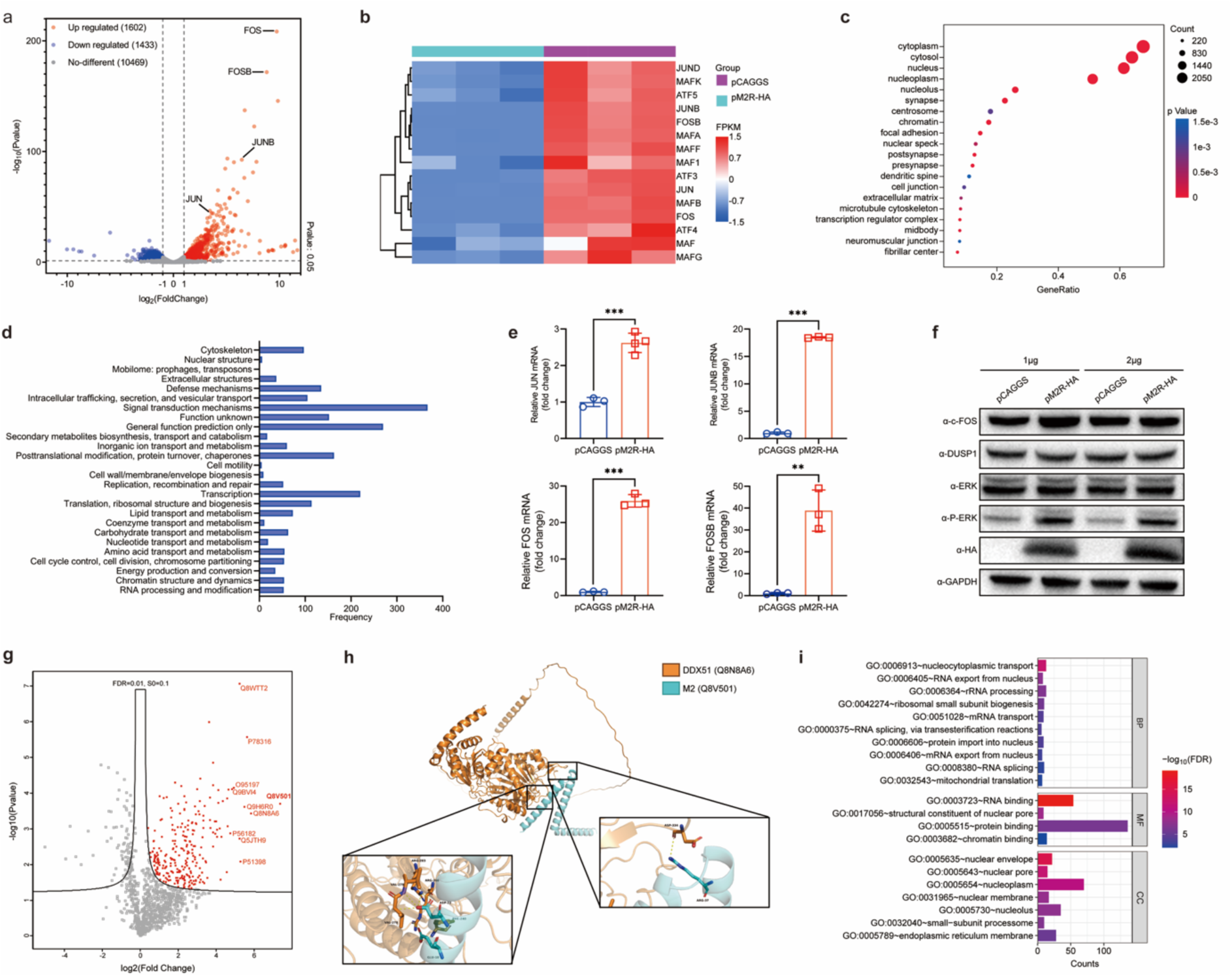
M2 modulates the ERK/AP-1 signaling pathway and activates associated transcription factors. The transcriptional impact of M2 was assessed via RNA-seq. HEK293T cells were transfected with either pCAGGS (vector control) or pM2R-HA for 24 hours, followed by mRNA extraction and RNA-seq analysis. **a** Volcano plot displaying differentially expressed genes in M2-expressing cells. Red dots represent upregulated genes; blue dots indicate downregulated genes. **b** Heatmap visualization of AP-1 family-related differentially expressed genes. Red and blue shades denote upregulation and downregulation, respectively, with darker colors indicating more significant changes. **c** GO cellular component enrichment analysis for differentially expressed genes. **d** KOG functional classification of differentially expressed genes in M2-transfected cells compared to control cells. **e** RT-qPCR analysis of AP-1 family factor transcription. HEK293T cells were transfected with pCAGGS or pM2R-HA and harvested at 24 hours post-transfection. mRNA levels were quantified by RT-qPCR. The data are shown as mean ± SD of three biological repeats; ***, P ≤ 0.001; **, P ≤ 0.01. **f** Altered protein expression of ERK/AP-1 pathway components. HEK293T cells were transfected with increasing concentrations of pM2R-HA or control pCAGGS for 24 hours. Whole-cell lysates were analyzed by western blotting. **g** The volcano plot displays the results of an IP-MS analysis identifying potential intracellular proteins interacting with M2, and proteins that fall within the defined threshold (FDR < 0.01 and log2 fold change > 0.1) are marked for further analysis. **h** AlphaFold3 predicts and analyzes the structural model of the interaction between M2 and DDX51. **i** The GO enrichment analysis of the target proteins identified through IP-MS.

**Fig. 7.**
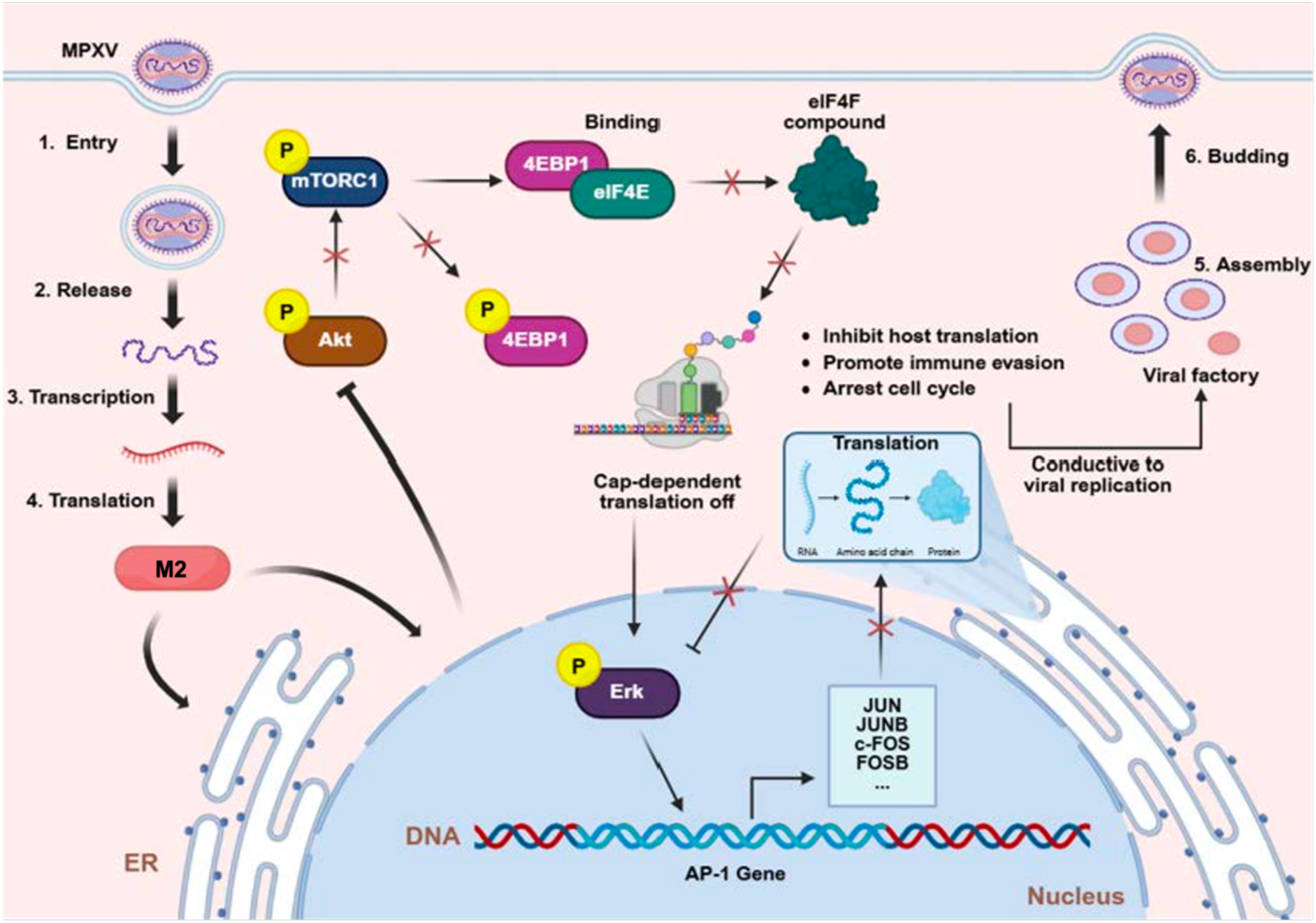
Signal transduction pathways for M2-triggered host shutdown. Upon viral entry, MPXV releases its genetic material into the host cell. The M2 gene is transcribed and translated in the cytoplasm, and the resulting protein localizes to the endoplasmic reticulum and nuclear envelope. Nuclear envelope-localized M2 inhibits nucleocytoplasmic transport, which subsequently suppresses Akt phosphorylation. This reduction leads to decreased phosphorylation of mTORC1. The inhibition of mTORC1 results in reduced phosphorylation of 4EBP1, leading to an increased association of hypophosphorylated 4EBP1 with eIF4E. This interaction impedes the assembly of the eIF4F complex, thereby suppressing cap-dependent translation. Additionally, impaired cap-dependent translation may enhance Erk phosphorylation, though the underlying mechanism remains unclear. Phosphorylated Erk translocate into the nucleus and activate transcription of AP-1 family genes (e.g., JUN, JUNB, FOS, FOSB). However, due to blocked nucleocytoplasmic transport, the resulting mRNAs cannot be exported efficiently for translation. Consequently, the negative feedback regulation of phosphorylated Erk by AP-1 proteins is disrupted, leading to sustained transcriptional activation of AP-1 target genes. Created with BioRender.com.

## Discussion

The intricate interplay between viruses and their hosts represents a continuous evolutionary arms race. Our study identifies the MPXV M2 protein could be a master orchestrator of host shutoff, leveraging a multi-pronged strategy that concurrently disrupts nucleocytoplasmic trafficking and inhibits translation initiation. While the VACV homolog L2 is traditionally characterized as an ER-resident protein essential for virion assembly [37, 41], M2 may have been evolved a distinct functional “exaptation.” By partially translocating to the nuclear envelope, M2 transcends its ancestral structural function, establishing a temporal and spatial switch during early infection potently contributes to virus-mediated host shutoff.

A defining feature of M2-mediated shutoff is the establishment of a “molecular double-lock” on host gene expression. First, M2 expression impairs nuclear import and mRNA export, potentially via interaction with the RNA helicase DDX51—a pivotal regulator of RNA splicing and export [44, 45] —thereby sequestering host poly(A)+ mRNAs within the nucleus and effectively decoupling transcription from translational machinery. While the directness of this effect and the molecular mechanism remain unclear, we cannot exclude indirect consequences of broader cellular perturbation. Second, M2 may impose a biochemical blockade in the cytoplasm by directly or indirectly antagonizing the Akt/mTOR/4EBP1 signaling axis. The consequent dephosphorylation of 4EBP1 promoyes sustained sequestration of eIF4E, dismantling the eIF4F complex required for cap-dependent translation [46, 47]. This synergistic redundancy ensures that even if host transcripts bypass the nuclear envelope, they encounter a paralyzed translational apparatus, rendering the host de novo proteome non-functional. However, the relative contributions of transport inhibition and translational suppression to the global host-shutoff phenotype remain to be determined, and whether these processes are functionally linked or independent awaits further investigation.

Perhaps the most intriguing finding is the “transcriptional trap” elicited by transfected M2. We observed a paradoxical upregulation of stress-responsive genes, particularly the AP-1 family (FOS, JUN), driven by sustained ERK phosphorylation. Under normal physiology, such a robust transcriptional burst would trigger an effective inflammatory response and subsequent negative feedback loops to resolve the signaling [48]. However, M2-induced nucleocytoplasmic blockade renders this mobilization futile. The host cell vigorously transcribes defense messages that fail to undergo normal translation, leading to a “silent” immune state at the protein level that may fuel a vicious cycle of resource exhaustion. This decoupling of transcription and translation represents may contribute to a sophisticated viral interference mechanism that enables unrestrained viral replication; however, whether M2 exerts a direct role during viral infection remains to be investigated.

The functional divergence between MPXV M2 and VACV L2 provides critical insights into poxvirus evolution and pathogenesis. Our virulence assays demonstrate that the acquisition of this host-shutoff capability significantly enhances viral fitness and pathogenicity, potentially contributing to the superior transmissibility observed in recent MPXV outbreaks. This “moonlighting” behavior, where a single protein executes both structural and immunomodulatory functions [49, 50], exemplifies the coding efficiency of large DNA viruses. From a translational perspective, the multifunctional nature of M2 identifies it as a vulnerable chokepoint for therapeutic intervention. Given the high conservation of its functional core across the Orthopoxvirus genus, disrupting M2’s interaction with the nuclear transport machinery or its suppression of the mTOR pathway could potentially restore host antiviral immunity. In conclusion, our investigation of the M2-dependent shutoff landscape not only advances the understanding of MPXV pathogenesis but also establishes a new paradigm for how DNA viruses orchestrate a comprehensive takeover of the host cell’s molecular flow.

This study has several caveats. First, preliminary screening experiments relied on transient M2 transfection in HEK293T cells, which may not fully recapitulate authentic MPXV infection, where M2 is co-expressed with other viral proteins at physiological levels. Overexpression may introduce artifacts stemming from protein mislocalization, perturbed molecular interactions, or competition for limited cellular resources. Second, our conclusions regarding nucleocytoplasmic transport and translational suppression remain largely correlative, with direct causal links yet to be established. Third, the M2–DDX51 interaction, identified via IP-MS and predicted by structural modelling, awaits biochemical validation. Fourth, polysome profiling data, while consistent with our model, are subject to technical limitations and require further refinement. Notwithstanding these caveats, integrating systematic functional screening with multiple experimental validations, including functional assays in viral infection models, yields preliminary insights into the potential role of MPXV M2 in host shutoff.

## Materials and Methods

### Cells and Reagents

In this work, HEK293T and Vero cells as well as BS-C-1 cells were cultivated in Dulbecco’s Modified Eagle Medium (DMEM, Gibco) supplemented with 10% (v/v) fetal bovine serum (FBS, Gibco) and 1% pen strep (Gibco). One day before each experiment, cells were seeded at an appropriate confluence in plates and incubated at 37°C overnight with 5% CO2. For DNA plasmid transfection, cells were seeded on plates 1 day before and transfected using Lipofectamine 3000 (Thermo) according to the manufacturer’s instructions.

### Construction of plasmids

The genes of the Monkeypox virus strain Zaire-96-I-16 (GenBank accession number NC_003310.1) were synthesized by GenScript and subcloned into pCAGGS vector with HA tag using FastDigest EcoRI (Thermo Fisher Scientific), FastDigest XhoI (Thermo Fisher Scientific), 2 × Phanta Flash Master Mix ClonExpress (Vazyme), SanPrep Column DNA Gel Extraction Kit (Sangon Biotech) and II One Step Cloning Kit (Vazyme). All primers used for molecular cloning were synthesized by TSINGKE gene factory. Plasmids were purified using Endo-Free Plasmid Midi Kit (OMEGA). All constructed plasmids were confirmed by sequencing, and no unexpected mutations occurred.

### Antibodies

The catalog numbers and dilution factors of the primary and secondary antibodies used in this study are illustrated in Table 1.

**Table 1.** The information of the antibodies used in this study.

| Antibody | Catalog no. | Clone name | Manufacturer |
| --- | --- | --- | --- |
| Anti-puromycin | MABE343 | Rabbit | Sigma |
| Anti-GAPDH | 60004-1 | Mouse | Proteintech |
| Anti-HA | 51064-2 | Rabbit | Proteintech |
| Anti-BIP | 11587-1 | Mouse | Proteintech |
| Anti-Lamin B | 12987-1 | Rabbit | Proteintech |
| Anti-Calnexin | C4731 | Rabbit | Sigma |
| Anti-Cas9 | 26758 | Rabbit | Proteintech |
| Anti-4EBP1 | 60246 | Mouse | Proteintech |
| Anti-P-4EBP1 | AP1363 | Rabbit | ABclone |
| Anti-mTOR | 7C10 | Rabbit | CST |
| Anti-P-mTOR | D9C2 | Rabbit | CST |
| Anti-EIF4G | 15704-1 | Rabbit | Proteintech |
| Anti-P70 (S6K) | 14485-1 | Rabbit | Proteintech |
| Anti-Akt | 10176-2 | Rabbit | Proteintech |
| Anti-P-Akt | 66444-1 | Mouse | Proteintech |
| Anti-Renilla | PM047 | Rabbit | MBL |
| Anti-c-FOS | 66590 | Mouse | Proteintech |
| Anti-DUSP1 | 49296 | Rabbit | SAB |
| Anti-ERK | A4782 | Rabbit | ABclone |
| Anti- P-ERK | AP0974 | Rabbit | ABclone |

### Renilla Luciferase Assay

The luciferase reporter plasmid pRL-SV40 used in this study was described previously [51]. Each preconstructed plasmid with virus functional gene and reporter plasmid pRL-SV40 were co-transfected into HEK-293T cells using Lipofectamine 3000 according to the manufacturer’s instructions. Cells were harvested after transfecting 24 hour, and a Renilla luciferase reporter assay system (Promega) was used to determine the luciferase activities of the lysed cells.

### Ribopuromycylation assay

Cells were infected or transfected for 24 hour and incubated with 3 μM of puromycin (MCE) in 10% FBS/DMEM for 1 hour. Cells were harvested in Laemmli sample buffer supplement with 2-mercaptoethanol and analyzed by immunoblotting. Translation was estimated using ImageJ and normalized to empty vector controls.

### Immunoblotting

Cells were lysed in Laemmli sample buffer and boiled. Cell lysates were separated by sodium dodecyl sulfate polyacrylamide gel electrophoresis with 4 to 20% gradient gel (GenScript) and transferred onto poly (vinylidene difluoride) membranes (Millipore). Membranes were blocked in 5% non-fat powdered milk/Tris-buffered saline with 0.1% Tween 20 (TBST) for 1 hour at RT, incubated with primary antibodies in 2% BSA/TBST overnight at 4 °C, incubated with secondary horseradish peroxidase-conjugated antibodies in 2% BSA/TBST for 1 h at RT, and analyzed by Tanon 4600 Imaging System (Tanon).

### Confocal Immunofluorescence Assay

Cells were washed with PBS and fixed with 4% paraformaldehyde in PBS for 15 min at RT. Cells were permeabilized with 0.1% Triton X-100 in PBS for 5 min at RT and blocked with 1% BSA in PBST (0.2% Tween-20 in PBS). Primary and secondary staining were performed in 1% BSA/PBST. Primary antibodies used were the same as used for immunoblotting. Secondary antibodies were the Alexa Fluor series from Thermo Fisher Scientific. Slides were mounted with ProLong Gold Anti-Fade Reagent (Thermo Fisher Scientific), imaged by Leica SP8 model confocal microscope, and analyzed using program LAS-X as well as ImageJ.

### Total RNA synthesis detection

Total cellular RNA synthesis was assayed using the BeyoClick™ EU-488 RNA Synthesis Detection Kit. HEK293T cells were seeded one day prior to transfection and cultured for 24 h post-transfection, then labelled with 1 mM EU at 37 °C for 2 h. Cells were fixed in 4% paraformaldehyde, washed with 3% BSA/PBS, permeabilized with 0.3% Triton X-100, and processed for click chemistry. After further washing, nuclei were counterstained with Hoechst, and fluorescence images were acquired using a fluorescence microscope.

### Recombinant virus construction

Recombinant virus VM2R was generated by homologous recombination. Briefly, the homologous recombination cassette was designed to contain 500 bp upstream and 1000 bp downstream flanking sequences of the VACV L2R gene. The cassette, comprising the PE/L promoter, EGFP, HA tag and MPXV M2R coding sequence between the two homology arms and subcloned into pBluescript IISK(+) to yield plasmid pVACV_M2R_EGFP. BHK-21 cells seeded in 6-well plates were infected with wild-type VACV strain WR at 0.01, 0.05 or 0.1 MOI, followed by transfection with pVACV_M2R_EGFP to mediate homologous recombination. Homologous recombination was preliminarily assessed by CPE and EGFP fluorescence. Recombinant VM2R was isolated by repeated fluorescent plaque picking and eight rounds of plaque purification using low-melting-point agarose overlays. Viral genomic DNA was extracted from purified VM2R and wild-type WR using the Tiangen DP315-F viral DNA/RNA extraction kit. Primers targeting EGFP-HA-M2R and HA-M2R were designed with Primer Premier 5.0 and synthesized by Tsingke Biotechnology. PCR and agarose gel electrophoresis were performed to validate the recombinant viral genome.

### Nucleus cytoplasm separation assay

Cytosolic and nuclear fractions were prepared from adherent HEK293T cells using the Minute™ Cytoplasmic and Nuclear Extraction Kit (Invent Biotechnologies, cat. no. SC-003). Near-confluent cell monolayers were washed with ice-cold PBS before lysis in cytoplasmic extraction buffer on ice. Following lysis, cell lysates were collected and vortexed vigorously. After centrifugation at 4 °C, the supernatant corresponding to the cytosolic fraction was transferred to a pre-chilled tube. The remaining nuclear pellet was resuspended in nuclear extraction buffer and processed with repeated vortex-ice incubation cycles. The nuclear lysate was then clarified via the kit-provided filter cartridge by high-speed centrifugation. Purified cytosolic and nuclear protein fractions were aliquoted and stored at −80 °C for subsequent assays.

### Electrophoretic mobility shift assay

EMSA was performed using 6.5% native PAGE gels, prepared from 30% acrylamide, 10× TBE, 50% glycerol, 10% APS and TEMED. Gels were pre-run in 0.25× TBE at 120 V for 1.5 h on ice. Serially diluted protein was incubated with 1 μM 5′-Cy5-labelled oligo(dT) or template mRNA in Interaction buffer I (final volume 10 μL) for 45–60 min at room temperature in the dark. Glycerol was added to 1% (v/v) final concentration before samples were loaded. Electrophoresis was carried out at 120 V on ice for 1 h in 0.5× TBE. Gels were imaged using a Typhoon FLA 9500 laser scanner.

### mRNA Stability Assay

HEK293T cells were transfected with 1 μg/mL pEGFP for 24h before transfected with 1 μg/mL pL2R-HA or with 1 ug/mL pCAGGS for 12h. Actinomycin D 5 μg/mL) were added to arrest transcription for various times. Total RNA was extracted using a TRIzol Plus RNA Purification Kit (Invitrogen) and analyses by RT-qPCR.

### RT-qPCR

Total RNA was extracted from cells using TRIzol Plus RNA Purification Kit (Invitrogen) according to the manufactures’ instructions. For one-step RT-qPCR, RNA was quantified using ChamQ Universal SYBR qPCR Master Mix (Vazyme). Analyses were performed using CFX Connect (BIO-RAD).

**Table 2.**
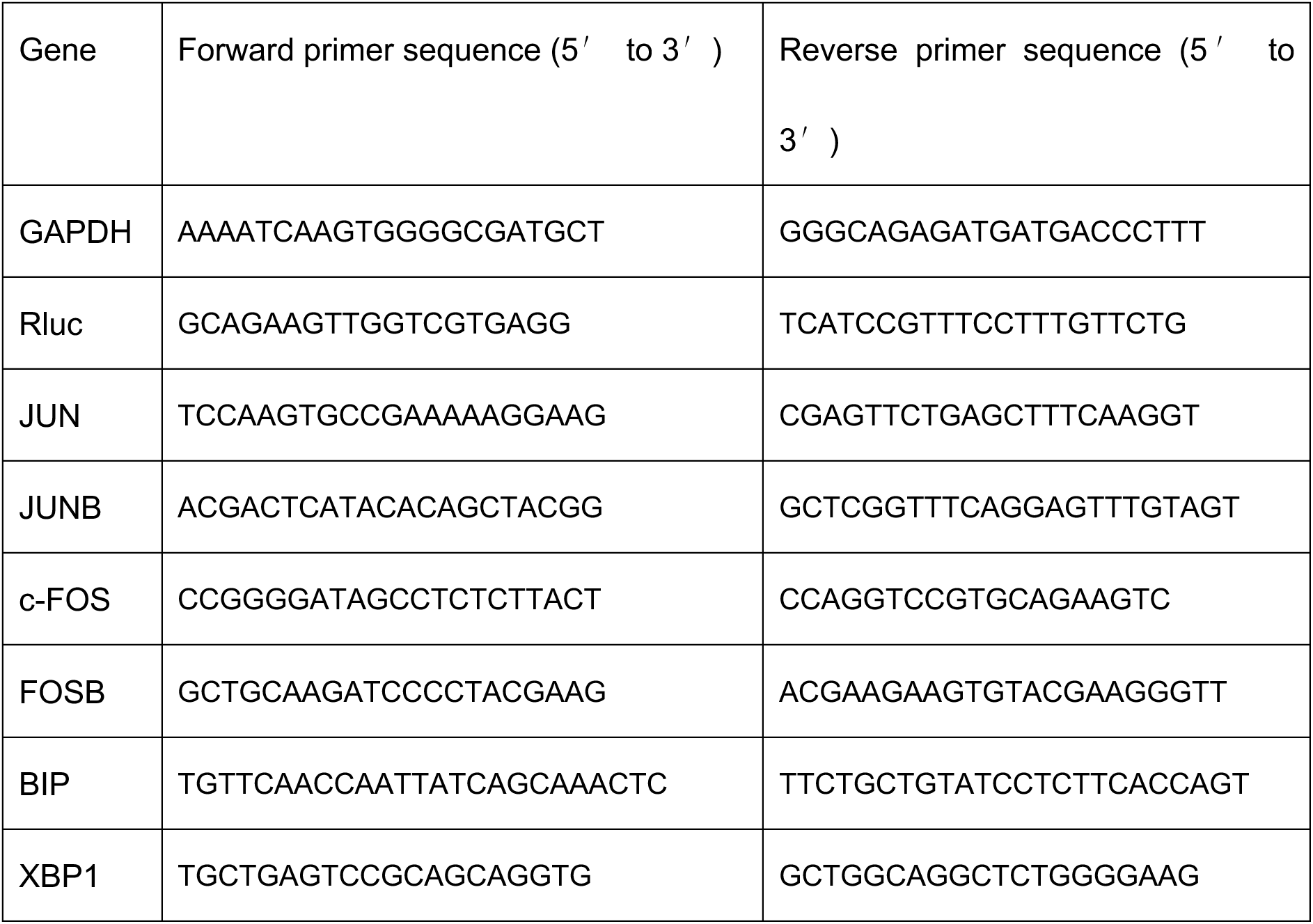
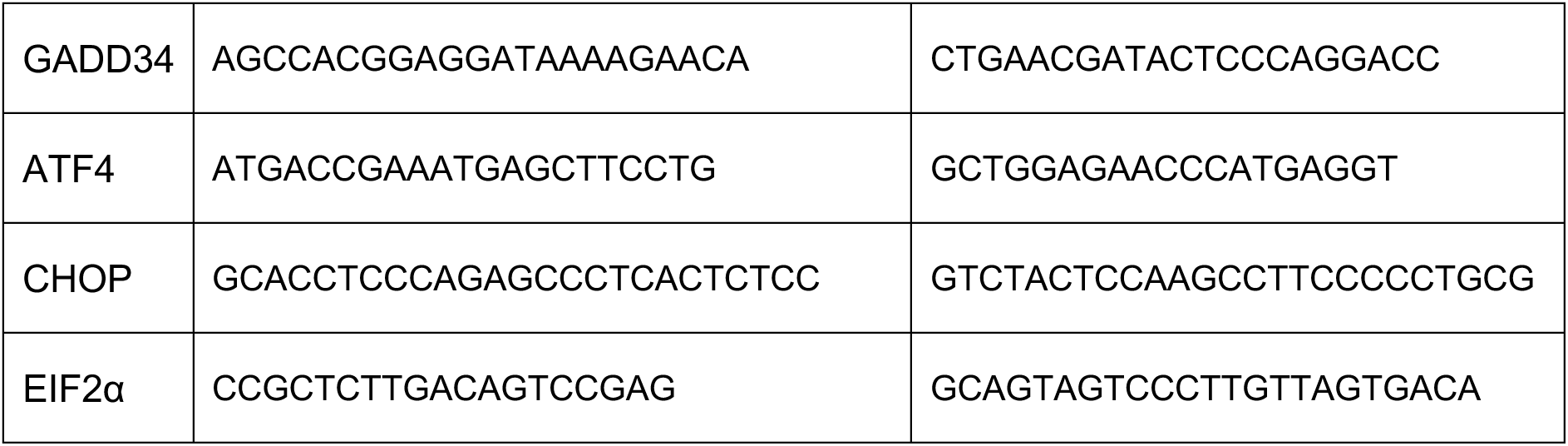
Sequences of primers used for qRT-PCR.

### Polysome Profiling

HEK293T cells were treated with 50 μg/mL CHX at 37°C for 10 min, washed with ice-cold PBS, incubated with 50 µg/mL CHX/PBS at 4 °C for 20 min, and lysed by adding polysome lysis buffer (200 mM KOAc, 50 mM Tris, pH 7.4, 10 mM MgCl2, 1 mM dithiothreitol, 0.5% IGEPAL CA-630, RNaseOUT, 50 µg/mL CHX and protease inhibitor mixture) on ice for 15 min. Cell lysates were centrifuged at 14,000 rpm for 15 min to remove cell debris. The resulting supernatants, containing the ribosomal complexes, were carefully layered onto pre-formed 10%-50% (w/v) linear sucrose density gradients (prepared in 200 mM KOAc, 50 mM Tris-HCl pH 7.4, 10 mM MgCl₂, and 50 µg/mL CHX). Gradients were ultracentrifuged in a Beckman Coulter SW41 Ti rotor at 35,000 rpm for 2.5 hours at 4°C to achieve resolution of the ribosomal species. Following centrifugation, gradients were fractionated using a Brandel BR-188 tube piercer and a BioLogic LP system (Bio-Rad), with continuous monitoring of absorbance at 254 nm. The resulting polysome profiles were analyzed to quantify the areas under the curve corresponding to sub-polysomal (40S, 60S, 80S) and polysomal (≥ 2 ribosomes per mRNA) fractions. The polysome-to-monosome (P/M) ratio was calculated for quantitative comparisons of translational efficiency between experimental groups.

### ER Stress and UPR

HEK293T cells in culture were transfected with 1 μg/mL pL2R-HA or with 1 μg/mL pCAGGS as well as treated with 1 μM TG (Sigma) as a positive control for UPR induction. BiP, XBP1, CHOP, ATF4, GADD34 and eIF2α cDNAs were quantified via RT-qPCR assay.

### Statistical analysis

Results from all studies were compared with unpaired two-tailed Student’s t test. P values less than 0.05 were considered significant.

## Acknowledgements

We are indebted to Prof. Cong Zeng at the Key Laboratory of Medical Molecular Virology (MOE/NHC/CAMS), School of Basic Medical Sciences, Fudan University, for his thoughtful guidance on partial experimental design. We also appreciate the Large Instruments Sharing Platform at the School of Basic Medical Sciences, Fudan University, for providing technical support and equipment access to confocal fluorescent microscopy and flow cytometer. This work was supported by grants from Shanghai Municipal Science and Technology Major Project (ZD2021CY001) and the National Natural Science Foundation of China (82030062 and 32571391).

## Contributions

Contribution: H.X. and L.W. conceived and designed the study. H.X. and Y.Y. performed the experiments and data analysis. W.W. and H.X. wrote the manuscript, and L.W. critically edited it. All authors reviewed and approved the manuscript.

## Extended Data figures

**Extended Data Fig. 1.**
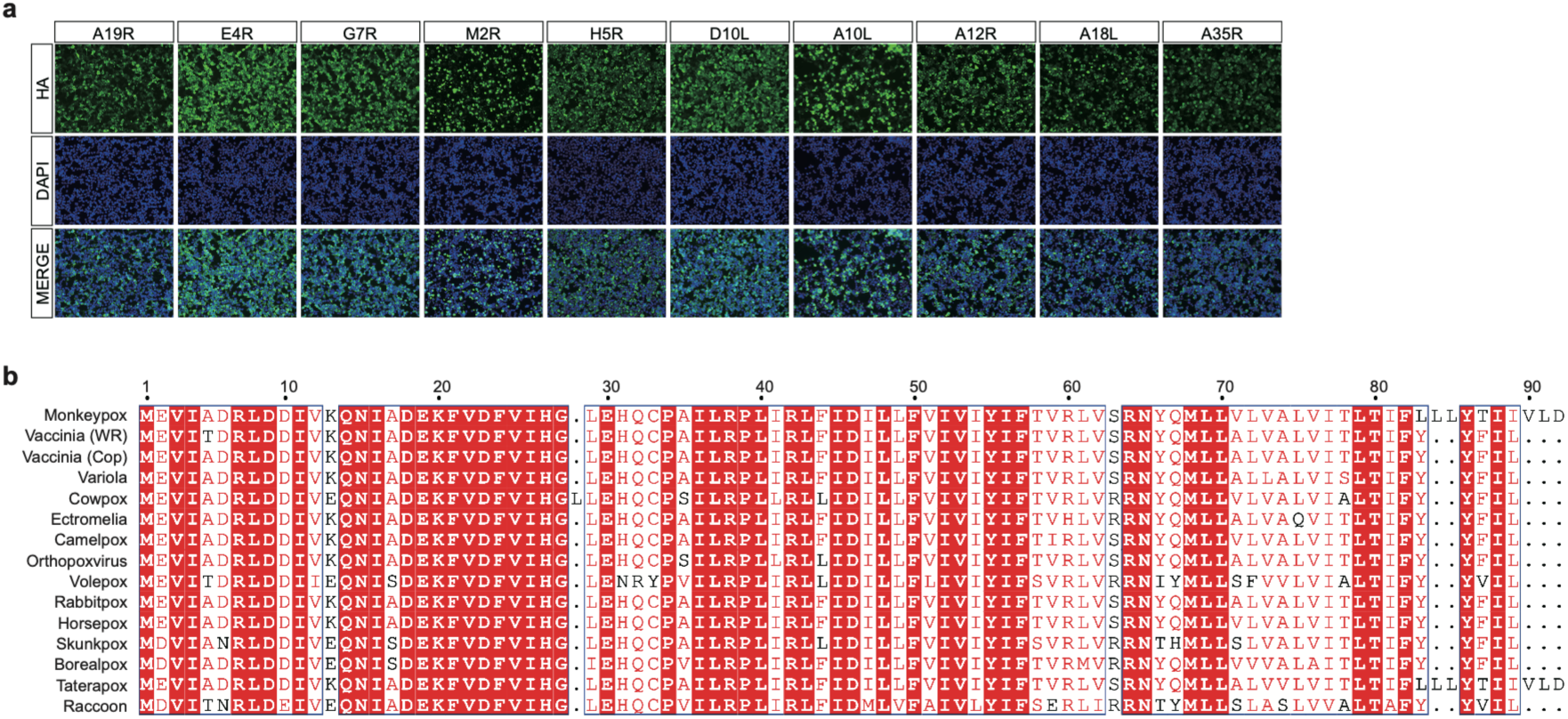
Expression and sequence conservation of poxviral M2R homologs. **a** Expression levels of plasmids encoding viral genes. Plasmids encoding MPXV viral proteins were transfected into HEK293T cells and assessed using immunofluorescence assay. **b** Amino acid sequence alignment of M2-related open reading frames from representative species of the poxvirus genus.

**Extended Data Fig. 2.**
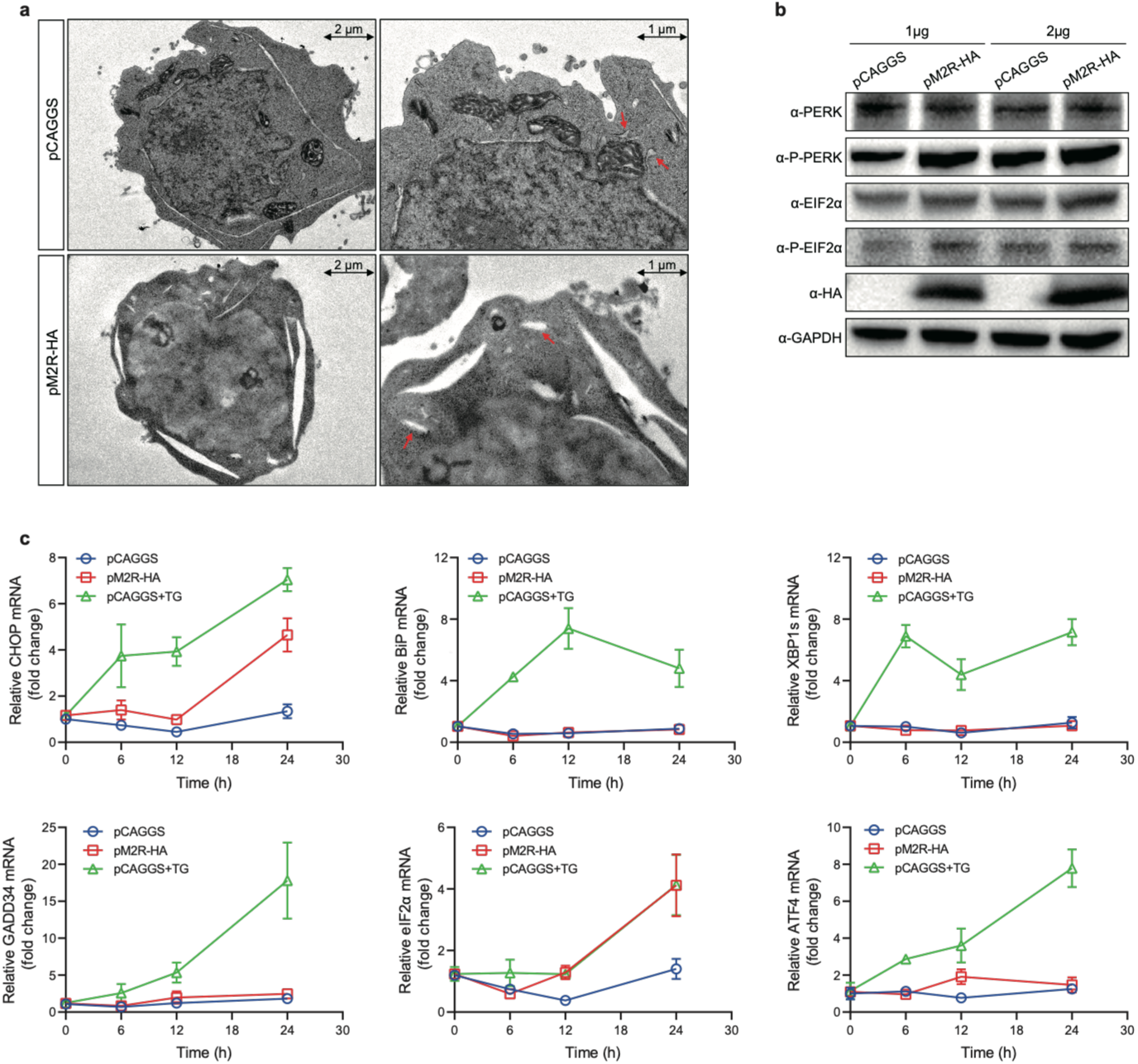
M2 and Endoplasmic Reticulum Stress. **a** M2-induced morphological changes in the endoplasmic reticulum. HEK293T cells were transfected with either pCAGGS or pM2R-HA, followed by fixation, embedding, and observation of ER morphology using transmission electron microscopy. **b** Altered protein levels of ER stress-related factors. HEK293T cells were transfected with increasing concentrations of pCAGGS or pM2R-HA. Whole-cell lysates were harvested 24 hours post-transfection and subjected to western blot analysis. **c** Transcriptional changes in ER stress-related factors. HEK293T cells were transfected with different concentrations of pCAGGS or pM2R-HA. At 12 hours post-transfection, thapsigargin (TG), an ER stress inducer, was added to pCAGGS-transfected cells. Cells were collected at the indicated time points and analyzed by RT-qPCR.

**Extended Data Fig. 3.**
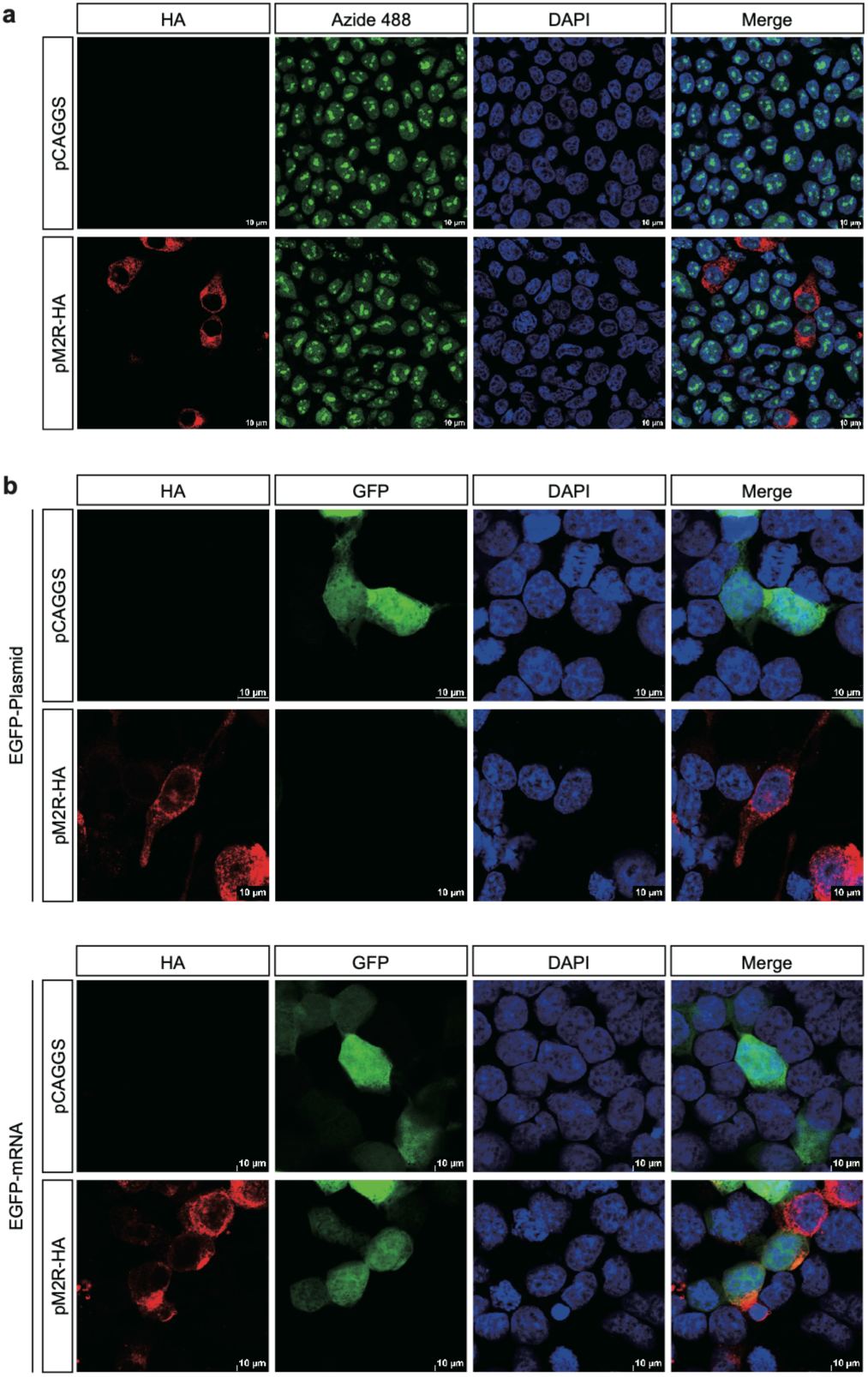
M2 does not suppress global transcription in host cells. **a** Examination of the impact of M2 on total RNA synthesis. HEK293T cells were transfected with pM2R-HA. At 24 hours post-transfection, newly synthesized mRNA was labeled using the uridine analog 5-ethynyl-uridine (EU), followed by click chemistry reaction with Azide 488 for fluorescence detection under microscopy. **b** Investigation of the effect of M2 on host transcriptional or translational processes. HEK293T cells were co-transfected with M2 and EGFP-expressing plasmid or mRNA. EGFP protein expression was observed using fluorescence microscopy.

**Extended Data Fig. 4.**
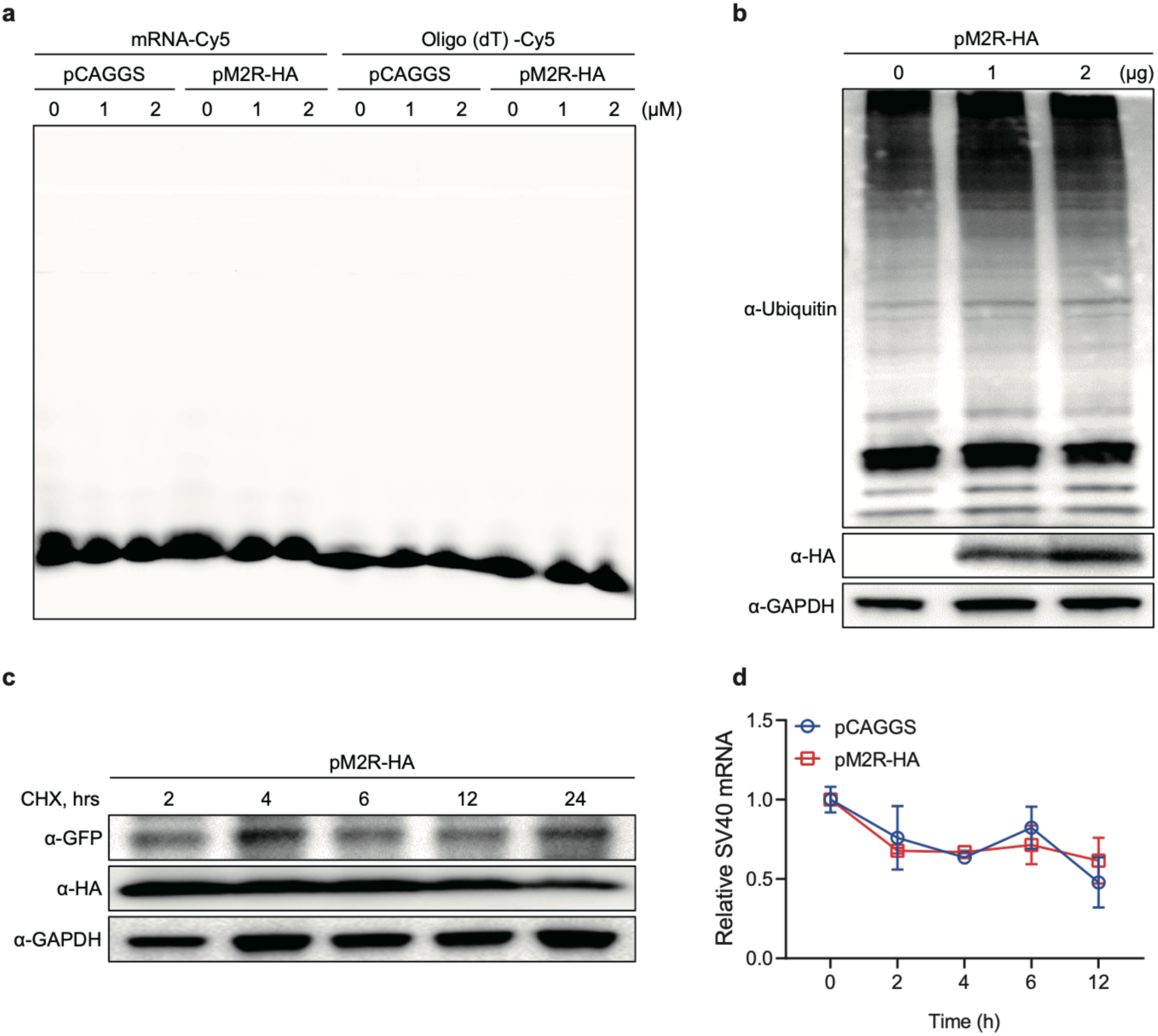
Effects of M2 on Intracellular Protein and mRNA Stability. **a** Assessment of M2 binding to mRNA and DNA. HEK293T cells were transfected with varying concentrations of pM2R-HA or empty vector. At 24 hours post-transfection, cell lysates were subjected to immunoprecipitation (IP) using HA-conjugated magnetic beads to pull down M2-HA. The potential binding ability of M2 to mRNA and DNA was analyzed using electrophoretic mobility shift assay (EMSA). **b** Effect of M2 on global ubiquitination levels. HEK293T cells were transfected with increasing concentrations of pM2R-HA. After 24 hours, whole-cell lysates were analyzed by western blot to detect changes in ubiquitinated proteins. **c** Effect of M2 on protein stability. HEK293T cells were co-transfected with pM2R-HA and a GFP-expressing plasmid. At 12 hours post-transfection, cycloheximide (CHX) was added to inhibit de novo protein synthesis. Cells were harvested at indicated time points, and GFP protein levels were analyzed by western blot. **d** Effect of M2 on mRNA stability. HEK293T cells were co-transfected with pM2R-HA and pRL-SV40. At 12 hours post-transfection, actinomycin D (ActD) was administered to block transcription. Cells were collected at specified time intervals, and SV40-derived mRNA levels were quantified using RT-qPCR.

**Extended Data Fig. 5.**
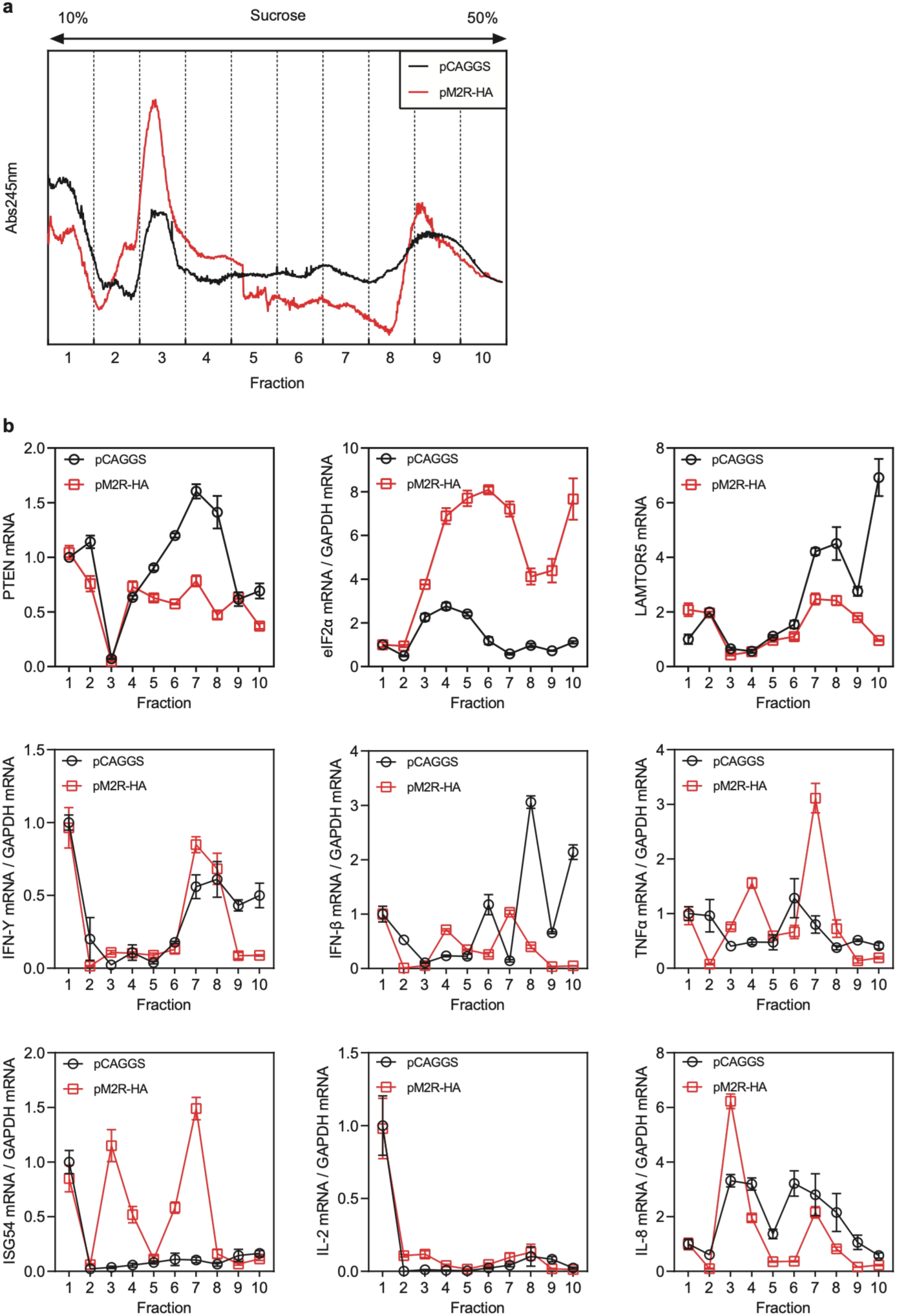
Analysis of M2 Effects on Ribosomal Translation. **a** Polysome profiling assay. HEK293T cells were transfected with either pCAGGS or pM2R-HA for 24 hours, followed by cell lysis and fractionation via sucrose gradient centrifugation. Polysome profiles were subsequently analyzed to evaluate ribosomal distribution. **b** mRNA analysis across ribosomal translation phases. mRNAs were extracted from individual fractions obtained by sucrose gradient centrifugation and subjected to RT-qPCR to quantify transcript distribution patterns.

